# MCM8/9 and FANCD2 interact within a shared pathway in response to replication stress caused by DNA crosslinks

**DOI:** 10.1101/2025.08.07.669127

**Authors:** Rashini Y. Baragama Arachchi, Desmond C. Okafor, Andrew J. Snyder, Michael A. Trakselis

## Abstract

The Fanconi anemia (FA) protein FANCD2, and MCM8/9 heterohexameric helicase complex are critical for maintaining genomic integrity in response to replication stress. However, the nature of their relationship remains unclear. Here, we show that MCM8/9 physically interacts and functionally cooperates with FANCD2 during the repair of DNA interstrand crosslinks (ICLs). Using immunofluorescence and co-immunoprecipitation studies, we show that MCM8/9 interacts with FANCD2 through its core domain, independently of DNA. FANCD2 is essential for the recruitment of MCM9 to ICL damage induced nuclear foci and acts downstream the FANCD2I monoubiquitination. Although MCM8/9 foci formation requires its intact ATPase activity, BRCv motif and HROB, these are not required for FANCD2 binding, highlighting a distinction between physical interaction and functional activation. Interestingly, FANCD2 foci formation increase in MCM8 or MCM9 knockout cells, suggesting that MCM8/9 functions to mitigate replication associate stress. γH2AX DNA damage assays and cell survival assays show that combined loss of MCM9 and FANCD2 do not cause any additive DNA damage beyond individual knockouts, indicating an epistatic relationship and suggests they function in the same DNA repair pathway. Together, our findings identify MCM8/9 as a downstream effector of the FA pathway critical for resolving ICL induced DNA damage.

**Highlights:**

- MCM8/9 interacts and colocalize with FANCD2 upon ICL DNA damage.
- FANCD2 is essential for recruitment of MCM9 to DNA damage site.
- MCM8/9 is epistatic to FANCD2 and within the same DNA damage response pathway.

## 1. Introduction

DNA replication stress causing replication fork stalling poses a significant threat to genomic stability, necessitating intricate cellular mechanisms to maintain genome integrity and suppress malignant transformation [1]. The Fanconi anemia (FA) pathway is a critical DNA damage response network, primarily orchestrating the repair of interstrand crosslinks (ICLs) and provides essential mitigation strategies for perturbed replication forks [2, 3]. Central to this pathway is FANCD2-FANCI (D2-I), a key DNA repair protein complex whose monoubiquitination functions as a key regulatory mechanism that coordinates multiple DNA repair processes to maintain replication fork stability [4]. There are at least 22 FA complementation group proteins, with D2/I functioning as a central player that integrates upstream signaling events with downstream repair activities [5]. Upon DNA damage or replication stress, the D2-I complex is monoubiquitinated (ubD2-I) by the FA core complex, which has an E3 ubiquitin ligase within one of the seven core proteins. ubD2-I forms a stable heterodimer on DNA that localizes within nuclear foci, where it coordinates key repair processes including ICL repair, translesion synthesis, and homologous recombination (HR), thereby protecting damaged or stalled replication forks [6, 7]. This post-translational modification is essential for D2-I’s function, as evidenced by the genomic instability observed in FA patients carrying mutations in FANCD2 or upstream FA genes [8, 9].

Recent advances in understanding replication fork progression have highlighted the importance of other DNA helicases, beyond the canonical heterohexameric MCM2-7 complex, that contribute to fork stability and progression. MCM8 and MCM9, are members of the same AAA+ minichromosome maintenance (MCM) family and have emerged as a specialized helicase complex that functions distinctly from MCM2-7 [10, 11]. Unlike their well-characterized counterparts, MCM8 and MCM9 form an alternating heterohexameric complex that exhibits unique biochemical properties and cellular functions [12-14]. MCM8/9 possesses 3’ to 5’ helicase activity and plays crucial roles in protecting stalled replication forks, HR mediated DNA repair for processing ICLs, and double strand breaks [15, 16]. Loss of MCM8 or MCM9 leads to reduced RAD51 recruitment, defective HR, and hypersensitivities to DNA-damaging agents such as mitomycin C (MMC), cisplatin, hydroxyurea (HU), and PARP inhibitors [13, 17, 18]. MCM8/9 requires binding of the activator protein, HROB, for proper recruitment and localization to damaged DNA sites [19, 20]. HROB interacts at an interface between MCM8 and 9 subunits to activate unwinding of various fork or HR substrates [21]. HROB-deficient cells phenocopy MCM8/9 knockout cells, showing impaired fork protection and reduced RAD51 accumulation [19]. Importantly, this MCM8/9/HROB complex appears to act within or around the FA pathway, where MCM8/9 is dependent for RAD51 (*i.e*. FANCR) foci and is counter to BRCA1 (*i.e*. FANCS) fork protection activity [13, 22], suggesting independent but coordinated roles in promoting genome integrity.

The phenotypic relationship between MCM8/9 and the FA pathway has only begun to converge through genetic and biochemical studies. Biallelic mutations in MCM8 and MCM9 have been identified in patients with premature ovarian insufficiency (POI) or failure (POF) and various chromosomal instability syndromes including cancer, phenotypes that overlap significantly with FA patients [23, 24]. Furthermore, cells deficient in MCM8 or MCM9 exhibit increased sensitivity to DNA crosslinking agents and display defects in homologous recombination that result in comparable radial chromosomes, suggesting functional integration with FA pathway components [15, 25]. However, the precise molecular mechanisms underlying MCM8/9 interaction with the FA pathway remain completely unresolved.

Here, we investigate the molecular and functional relationship between the MCM8/9 helicase complex and the FA pathway focusing on the intersection and interaction with FANCD2. Through a combination of immunofluorescence microscopy, cellular biology interactions, and cellular survival assays, we demonstrate direct physical and functional interactions between these critical genome maintenance proteins. Our findings reveal that MCM8/9 and FANCD2 exhibit reciprocal dependencies for proper crosslink dependent nuclear focus formation, establish direct protein-protein interactions, and coordinately regulate cellular responses to DNA damage. These results provide new insights into MCM8/9/HROB-FANCD2/I nexus and establish a direct molecular link between the MCM8/9/HROB helicase complex and the FA pathway. Our findings advance the understanding and intersection of the molecular networks that safeguard genomic stability.

## 2. Materials and methods

### 2.1 Cell culture

Wild type 293T, MCM8^KO^ and MCM9^KO^ in parental 293T cells [22] were cultured in Dulbecco’s Modified Eagle Medium (DMEM) (Gibco, Waltham, MA) supplemented with 10% fetal bovine serum (FBS) (Gibco) and 1% penicillin-streptomycin (Gibco) at 37 °C in a humidified incubator with 5% CO_²_.

### 2.2 Cloning

Truncated human MCM9 constructs were generated by PCR amplification from full-length MCM9 cDNA using Platinum^™^ SuperFi^™^ II DNA Polymerase (Fisher Scientific, Waltham, MA). PCR amplicons encoding amino acid residues 1–605, 1–648,1–800 and 605-1143 were cloned into the pEGFP-C2 vector (Clontech, San Jose, CA) at the multiple cloning site using a strategy incorporating a *StuI* restriction site (AGGCCT) for directional cloning (**Supplementary Table S1**). Construct-specific reverse primers were designed to encode the desired truncation points while incorporating a SV40 nuclear localization signal (NLS; PKKKRKV) to ensure efficient nuclear import. All plasmids were verified by sequencing (Plasmidsaurus, South San Francisco, CA) to confirm the correct sequence.

### 2.3 Transfection

293T WT, MCM9KO or FANCD2KO cells were transfected with pEGFPc2-MCM9^FL^, pEGFPc2-MCM9^605^, pEGFPc2-MCM9^648^, pEGFPc2-MCM9^800^, MCM9^605–1143^ (CTE), pEGFPC2-MCM9^FR687/8AA^ (BRCv^−^), pEGFPC2-MCM9^K358A^ (WA^-^) [22], pEGFPC2 (empty), pRPmCherry-hFANCD2 (VectorBuilder, Chicago, IL) plasmids using LPEI transfection reagent (Fisher Scientific) according to the manufacturer’s guidelines. Briefly, plasmid DNA and LPEI were mixed (1:3 ratio) and incubated at room temperature for 20 minutes before being added to cells in reduced serum Opti-MEM media (Gibco). For immunofluorescence experiments, six hours after transfection, the transfection medium was replaced with complete DMEM, and cells were incubated overnight. The following day, cells were treated with 3 µM mitomycin C (MMC) (Cell Signaling Technology, Danvers, MA) for 6 hours or 3 mM hydroxyurea (HU) (Fisher Scientific) for 5 hours, as indicated. Treated cells were then processed for immunofluorescence analysis.

For co-immunoprecipitation (co-IP) and immunoblotting experiments, cells were transfected with the indicated plasmids and incubated for 48 hours. The culture medium was then refreshed with complete DMEM, followed by treatment with 0.5 µM MMC or 2 mM HU overnight. Cells were subsequently harvested for co-IP or western blot analysis.

### 2.4 Immunofluorescence (IF)

Cells were seeded onto glass coverslips coated with poly-L-lysine (Newcomer Supply, Waunakee, WI) and treated as indicated. Following treatment, cells were fixed in 4% paraformaldehyde for 15 minutes at room temperature, permeabilized with 0.1% Triton X-100 for 10 minutes, and blocked with 2.5% fish gelatin in PBS at 4 °C overnight. Coverslips were incubated with primary antibodies α-FANCD2 (Santa Cruz, sc-2002) (1:500) or α-γH2A.X (Abcam, ab26350) (1:500) for 1 h at 37 °C in a humidified chamber. All antibody dilutions were prepared in 2.5% BSA in PBST. Cells were washed three times in PBST and incubated with 1:1000 dilution of the α-mouse Cy3 (ab97035, Abcam, Cambridge UK) secondary antibody for 30 min at 37 °C in a humidified chamber. Coverslips were then washed three times with PBST. Cells were mounted in mounting media containing DAPI (ProLong Diamond Antifade Mountant with DAPI, Thermo Fisher) and sealed with clear nail polish and imaged under a FV-1000 epifluorescence or FV-3000 confocal laser scanning microscope (Olympus Corp., Center Valley, PA). Images were processed with Fluoview (Olympus, v.4.2b) or cellSens software (Olympus, v2). FANCD2, MCM9 or γH2A.X foci counts from epifluorescence images were analyzed using ImageJ [26]. The colocalization of representative signals in different channels was examined using Fiji (v.1.54p) with the Coloc 2 plugin using Mander’s overlap coefficient (MOC) as it normalizes signal intensities between components [27, 28]. Data were analyzed for any statistically significant differences using GraphPad Prism (v. 10.4, San Diego, CA).

### 2.5 Co-Immunoprecipitation (co-IP)

Co-immunoprecipitation was performed using the ChromoTek GFP-Trap (for GFP-tagged constructs) (Proteintech, Rosemont, IL) or the Thermo Scientific Pierce Co-IP Kit (for endogenous proteins), following the manufacturers’ protocols. Briefly, cell lysates were prepared in ice cold lysis buffer (0.025 M Tris, 0.15 M NaCl, 0.001 M EDTA, 1% NP-40, 5% glycerol; pH 7.4) supplemented with protease/phosphatase inhibitors (Sigma-Aldrich), DNase 1 (final concentration 150 U/mL) (Fisher Scientific) and MgCl2 (final concentration 2.5 mM). Total protein concentrations were evaluated using BCA assay (Pierce BCA Protein Assay kit, Fisher Scientific). Equal concentrations of lysates were incubated with GFP-Trap beads (1 hour at 4°C with continuous mixing) or with antibody-crosslinked resin (overnight at 4°C with continuous mixing). Beads were washed four times with washing buffer provided in the respective kits, and bound proteins were eluted in 2x SDS-sample loading buffer for subsequent immunoblot analysis.

### 2.6 Immunoblotting

Equal amounts of protein (50 μg) were loaded into 8% SDS gels or 4-10% bis tris gradient NuPAGE gels (Fisher Scientific) and separated by polyacrylamide gel electrophoresis (PAGE). Proteins were transferred onto PVDF or nitrocellulose membranes (Cytiva, Marlborough, MA). Membranes were blocked in 5% non-fat dry milk in TBST (10 mM Tris-HCl [pH 7.4], 150 mM NaCl and 0.1% Tween-20) and probed with the indicated primary antibodies overnight at 4 °C with gently rocking. Membranes were washed 3 times with TBST and probed with HRP-conjugated secondary antibodies, α-mouse HRP (Novex A16072, Barberton, OH) or α-rabbit HRP (Novex A16096) for 1 hour at RT. The blots were washed three times with TBST and developed using chemiluminescence (sc-2048, Santa Cruz, Dallas, TX) and imaged with a ChemiDoc Imaging System (Bio-Rad, Hercules, CA).

### 2.7 Cell synchronization by double thymidine block

293T, MCM8^KO^ and MCM9^KO^ cells were synchronized at the G1/S boundary using the standard double thymidine block protocol. Briefly, cells were seeded in 10 cm^²^ dishes and grown in DMEM supplemented with 10% fetal bovine serum (FBS) until 40% confluency. To initiate synchronization, 10 mM thymidine (TCI America, Portland, OR) was added to the culture medium, and cells were incubated at 37 °C in a humidified incubator with 5% CO_2_ for 18 hours. Following the first thymidine block, the medium was aspirated, and cells were washed three times with pre-warmed phosphate-buffered saline (PBS). Cells were then released into the cell cycle by replacing the medium with fresh DMEM containing 10% FBS (without thymidine) and incubated for 8 hours. A second thymidine block was applied by reintroducing DMEM supplemented with 10% FBS and 10 mM thymidine for another 18 hours. Finally, cells were washed three times with warm PBS and released into fresh DMEM with 10% FBS. Cells were either treated with MMC or left untreated as indicated and processed for immunoblotting.

### 2.8 Generation of FANCD2 knockout cells

FANCD2 knockout (FD2KO) in 293T parental cells was generated using CRISPR-Cas9 gene targeting. Guide RNAs (gRNAs) (**Supplementary Table S1**) targeting exon 11 of the FANCD2 gene were selected based on sequences previously published [29] and cloned into the pSpCas9(BB)-2A-Puro (PX459) V2.0 vector (Addgene #62988) using *BbsI* restriction sites. HEK293T cells were transfected with this recombinant plasmid using LPEI (Fisher Scientific). After 48 hours, cells were transferred to selection media containing 1 μg/ml puromycin (Fisher Scientific) and cultured for an additional 72 hours with fresh puromycin containing media replaced daily to maintain selection and promote enrichment of transfected populations. Single cells were sorted into 96-well plates using a FACS Melody cell sorter (BD Biosciences, Franklin Lakes, NJ). To promote clonal expansion, single cells were cultured in complete DMEM supplemented with 30% filtered (0.22 μm membrane filter) conditioned media, prepared from a day-old HEK293T supernatant. Established clones were screened for successful FANCD2 knockout by immunoblotting and immunofluorescence to confirm the absence of FANCD2 protein expression.

### 2.9 siRNA mediated gene silencing

Multiple siRNA duplex oligonucleotides targeting FANCD2, MCM9, HROB, and non-target control (**Supplementary Table S1**) MISSION ® predesigned siRNAs (Sigma-Aldrich, St. Louis, MO) were transfected at a final concentration of 25 nM using Dharmafect1 (Dharmacon, Lafayette, CO) according to the manufacturer’s instructions. A second transfection was performed 48 hours after the first, and cells were incubated with siNRA complexes for a total of 72 hours post-initial transfection. After 72 hours of incubation with siRNA, media was replaced with fresh complete DMEM, treated as indicated, and cells were harvested for immunofluorescence or immunoblotting.

### 2.10 Short-term viability assays

Cell viability of single gene depleted MCM9KO, FANCD2KO or co depleted (MCM9KO: siFANCD2 or FANCD2KO: siMCM9) or WT 293T was assessed using CellTiter-Blue cell viability assay kit (Promega, Madison, WI) following the manufacturer’s instructions. Briefly, single gene depleted or co depleted cells (5000 cells/well) were seeded in 96-well plates, treated with a range of MMC concentrations as indicated for overnight (18 hours). Four hours prior to the end of MMC treatment, CelTiter-Blue (diluted in Ca^2+^/Mg^2+^ free PBS at 1:1 ratio) was added (20 μL/well) and plates were incubated for 4 hours at 37 °C in a humidified incubator with 5% CO_2_. Following incubation, fluorescence was measured at 560/590 nm using a microplate reader (Thermo Varioskan LUX Multimode Microplate Reader). Fluorescence data were normalized to untreated controls and data were analyzed using GraphPad Prism (v. 10.4).

### 2.11 Clonogenic survival assay

For colony formation assays, single gene depleted MCM9KO, FANCD2KO or co depleted (MCM9KO: siFANCD2, FANCD2KO: siMCM9) or WT HEK293T cells were seeded at low density (1000 cells/well) in 6-well plates. Twenty-four hours after seeding, cells were treated with the indicated concentrations of MMC and incubated for 24 hours. Following treatment, media was replaced with fresh complete DMEM media, and cells were allowed to grow to colonies for 10 -14 days. Colonies were washed with PBS, stained with 0.5% crystal violet, and counted using Fiji software’s ColonyArea plugin [30]. Survival fractions were calculated relative to untreated controls. Data was analyzed using GraphPad Prism (v. 10.4).

## 3. Results

### 3.1 MCM8/9 colocalizes and interacts with FANCD2

As we have previously shown that MCM8/9 partially colocalize with RAD51 and are required for RAD51 nuclear foci formation [22], we sought to look upstream in the crosslinking fork stalling protection pathway by examining any connection of MCM8/9 with D2-I. D2-I is an early effector within a fork protection pathway responding to crosslink damage that recruits proteins to stabilize the fork, including RAD51 and BRCA1 [31-33]. Amazingly, transfection of full length GFP-MCM9 results in MMC dependent nuclear foci that nearly completely colocalize with native FANCD2 (**Fig. 1A**). The Mander’s M2 overlap coefficient (MOC) for FANCD2 with GFP-MCM9 is greater than 0.9, measured for multiple cells. Neither GFP-MCM9 or FANCD2 form any significant foci in the absence of damage. As MCM8 forms a complex with MCM9, fluorescent colocalization experiments with FANCD2 were repeated with GFP-MCM8 with similar results (**Supplementary Figure S1A**).

**Figure 1.**
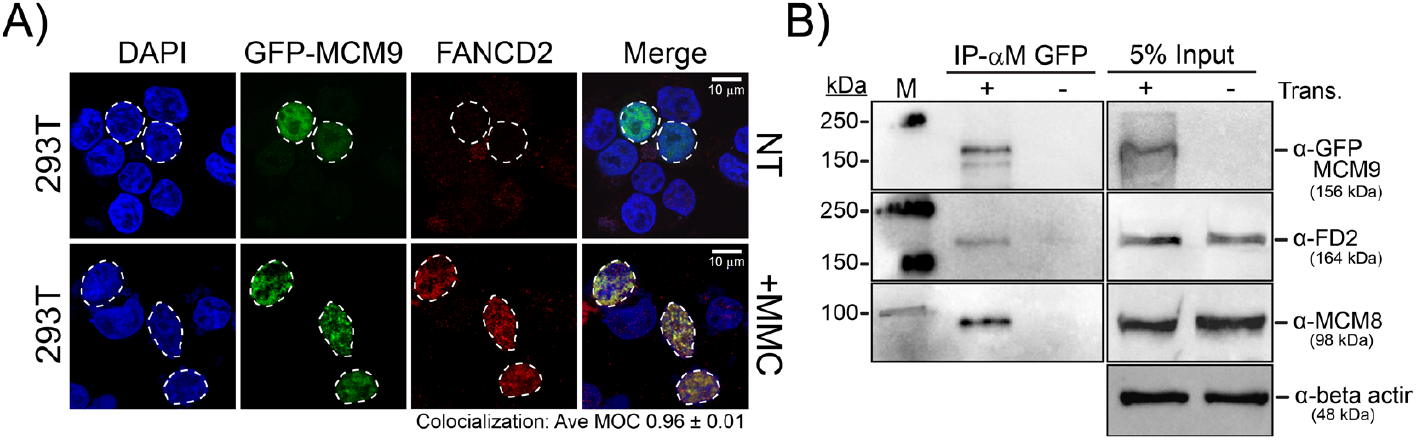
MCM9 colocalizes and interacts with FANCD2. A) Immunofluorescence of 293T cells transfected with pEGFP-MCM9 either nontreated (NT) or treated with 3 μM MMC (+MMC) for 6 hours. Endogenous FANCD2 (red) was stained with α-FANCD2 antibody. GFP-transfected cell nuclei are outlined (white dashed circle). The merge image shows near complete colocalization of MCM9 and FANCD2 foci treated with MMC. The average Mander’s M2 overlap coefficient (MOC) for FANCD2 overlapping with MCM9 is indicated. B) Co-IP of GPF-MCM9 with FANCD2 and MCM8. 293T cells were transfected (Trans.) with pEGFP-MCM9 or untransfected (- control) for 48 hours before treating with *0.5* μM MMC overnight. Whole cell lysates were IP’d with ChromoTek GFP-trap beads. Elutes were resolved by SDS-PAGE and immunoblotted for the indicated proteins. Left panel: IP’d; Right panel: 5% inputs. Molecular weight markers are indicated on the left and the position of constructs on the right with indicated theoretical molecular weights (kDa).

To determine whether MCM8/9 directly interacts with FANCD2 and does not just colocalize, various co-immunoprecipitation (co-IP) experiments were performed. Endogenous MCM9 co-IPs with both FANCD2 and MCM8 in the absence of DNA (+DNase I), and in reverse experiments, FANCD2 co-IPs with MCM8 (**Supplementary Figure S1B-C**). Further co-IP with GPF-Trap beads were performed with GFP-tagged MCM9 constructs with similar positive results for interacting with FANCD2 and MCM8 (**Fig. 1B**). Mutation of the Walker A – ATPase site (WA^-^) or the BRCv - BRC variant motif (BRCv^-^) in MCM9 retained the ability to co-IP with FANCD2 in the absence (NT) or presence of the fork stalling agents, hydroxyurea (HU) or MMC (**Supplementary Figure S2A-B**), suggesting that physical MCM9-FANCD2 interaction does not require these functional motifs. Therefore, neither DNA binding, the ATPase activity of MCM9, nor the recruitment of RAD51 is dependent on this interaction. The WA^-^ or BRCv^-^ mutations in MCM9 had been shown previously to not form MMC induced nuclear foci [13, 22]; however, these mutations do not affect the ability of FANCD2 to form foci with MMC (**Supplementary Figure S2C**). This demonstrates that MCM9’s ability to form functional nuclear foci requires its ATPase and BRC activities independently of its capacity to interact with FANCD2.

To further narrow down the site of FANCD2 interaction with MCM8/9, we truncated MCM9 at residues 605, 648, or 800 to keep the core DNA binding and ATPase domains while truncating past either the winged-helix (WH) domain or BRCv motif, respectively, but eliminate the long C-terminal extension (CTE) in MCM9 (**Fig.2A**). All MCM9 constructs containing the NTD and AAA+ domains are able to form a heterohexamer with MCM8 [14]. Interestingly, all constructs retained the interaction with FANCD2 through Co-IP (**Fig. 2B-C**), while GFP alone and the MCM9 CTE (605-1143) were unable to IP FANCD2 (**Supplementary Figure S3**), suggesting that the FANCD2 interaction occurs within the core hexameric structure. However, only GFP-MCM9^800^ was able to form MMC induced nuclear foci that colocalized with FANCD2 (**Fig. 2D**), confirming that the BRCv motif in MCM9 is still required to form MMC induced nuclear MCM9 foci formation, despite being dispensable for a physical interaction with FANCD2. To confirm that the various GFP-MCM9 deletion constructs can also restore the FANCD2 interaction in MCM9 knockout (9KO) cells (**Supplementary Figure S4**), we again performed Co-IP experiments and showed that all constructs still interact with FANCD2 and MCM8 (**Supplementary Figure S5**). Similar results are seen for colocalization with FANCD2 in 9KO cells, where only GFP-MCM9^FL^ restores complete colocalization and GFP-MCM9^800^ restores partial colocalization with FANCD2 (**Supplementary Figure S5C**).

**Figure 2.**
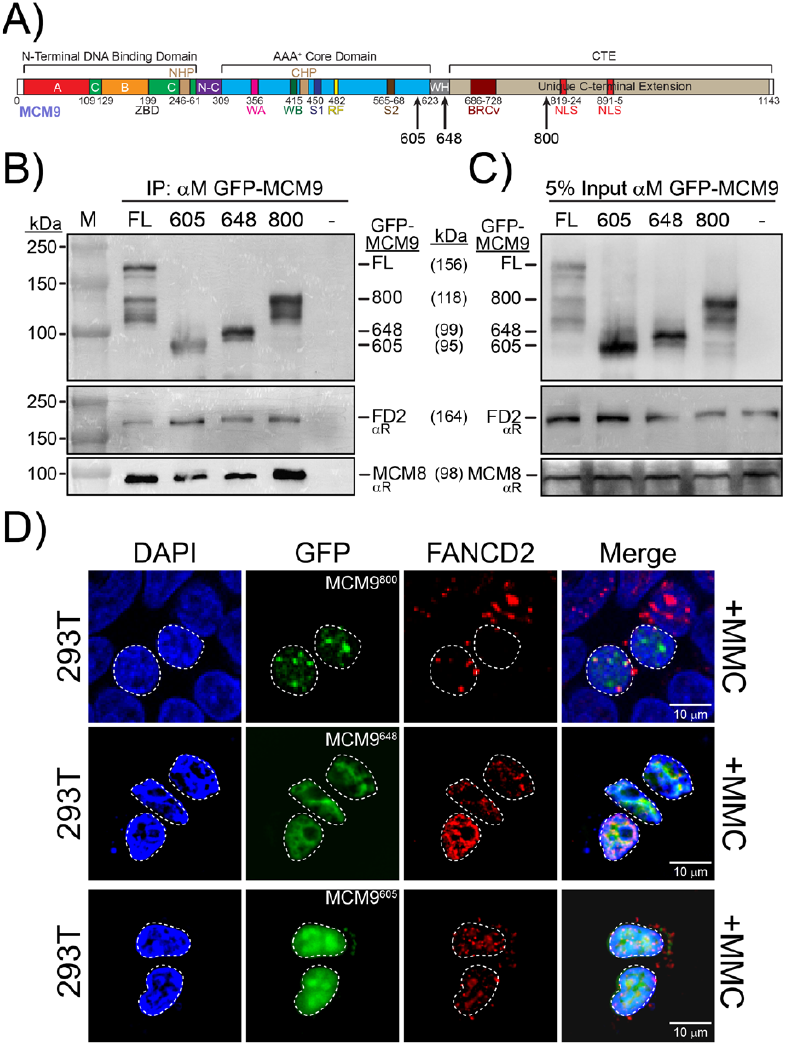
FANCD2 interaction occurs within the core domains of MCM8/9. A) Schematic of the MCM9 protein domain and motif organization and identifying the location of the different constructs of MCM9 used. B) 293T cells were transfected with pEGFP-MCM9^FL^, MCM9^605^, MCM9^648^, MCM9^800^ or untransfected (- control) for 48 hours before treating with 0.5 μM MMC overnight. Whole cell lysates were IP’d with ChromoTek GFP-trap beads. Elutes were resolved by SDS-PAGE and immunoblotted for the indicated proteins. C) 5% Input blots of the protein lysates. Molecular weight markers are indicated on the left and the position of constructs on the right with theoretical molecular weights (kDa). D) WT 293T cells were transfected with pEGF-MCM9 protein constructs (green) and treated with 3 μM MMC for 6 hours. Endogenous FANCD2 (red) was stained with α- FANCD2 antibody. GFP-transfected cell nuclei are outlined (white dashed circle).

The homologous recombination OB-fold protein (HROB) has been shown to interact with and stimulate MCM8/9 unwinding previously [19, 21]. To determine whether HROB is required for the interaction of GFP-MCM9 and FANCD2, we performed co-IP experiments where HROB was knocked down by siRNA. The presence of HROB was unnecessary for the MCM8-9-D2 complex formation by co-IP (**Fig. 3A-B**); however, siHROB eliminated the ability for GFP-MCM9 to form foci in the presence of MMC (**Fig. 3C**). Furthermore, siHROB did not affect FANCD2 foci formation.These results demonstrate that although MCM8/9 can physically interact with FANCD2, it is insufficient for functional colocalization at sites of replication stress. In addition to FANCD2, MCM8/9 foci formation also requires ATPase coupled and HROB stimulated unwinding activity as well as the BRCv motif to recruit RAD5, while FANCD2 foci formation occurs independently of MCM8/9 functional status. Therefore, it appears the MCM8/9 interacts with FANCD2 both on and off DNA and in the absence and presence of crosslink damage. However, MCM8/9 foci formation is somewhat independent of the FANCD2 interaction and instead depends more on treatment with MMC.

**Figure 3.**
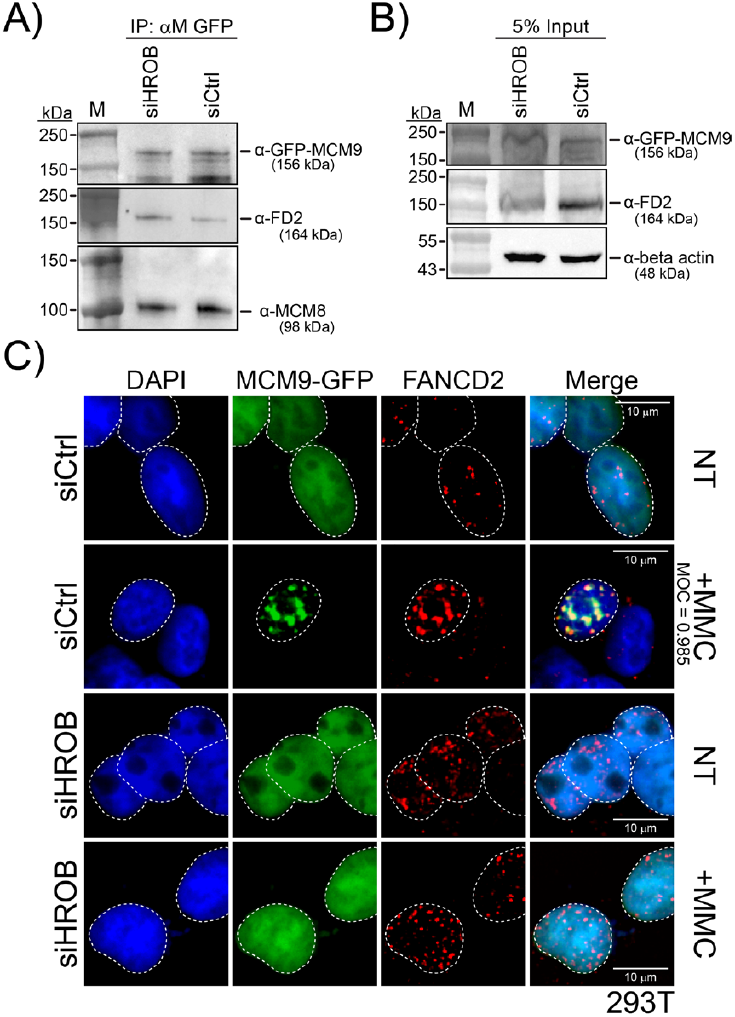
siRNA mediated depletion of HROB does not interrupt the MCM9-FANCD2 interaction. A) ChromoTek GFP-trap Co-IP or B) lysate 5% input of GFP-MCM9^FL^ with FANCD2 in siHROB depleted vs. universal non-target control (siCtl1) 293T cells treated with 0.5 μM MMC overnight. Molecular weight markers are indicated on the left and the position of constructs are on the right with theoretical molecular weights (kDa). C) Transfected and nontreated (NT) or MMC treated (+MMC) cells were imaged for GFP-MCM9 or FANCD2 foci and colocalization. Endogenous FANCD2 was stained with α-FANCD2 antibody. GFP-transfected cells are outlined (white dashed circle). The merge +MMC (siCrtl) image shows near complete colocalization of MCM9 and FANCD2 foci treated with MMC. MOC for FANCD2 overlapping with MCM9 is indicated.

### 3.2 Knockout of MCM8 or MCM9 increases FANCD2 foci

As the truncated or mutant constructs of MCM9 did not appear to affect the ability of MMC treated cells to form FANCD2 foci, we wanted to quantify whether the number of FANCD2 foci change when MCM8 (8KO) or MCM9 (9KO) are knocked out in 293T cells. Native FANCD2 foci were quantified in untreated (**Fig. 4A-B**) or MMC treated (**Fig. 4C**) cells. FANCD2 foci are visibly increased in the nuclei of untreated 8KO or 9KO cells, and when quantified, the number of FANCD2 foci increases significantly from an average of 2.4 ± 0.3 to 13.9 ± 0.8 or 13.3 ± 0.6, respectively, foci per cell. The increase in FANCD2 foci in the absence of MCM8/9 indicates that MCM8/9 are preventing a more severe DNA damage response by D2/I even in the absence of exogenous damage. FANCD2 foci increased even more with MMC treatment in WT (9.2 ± 0.3), 8KO (17.4 ± 1.0), or 9KO (14.6 ± 0.9) cells, as expected, and the transfection of GFP-MCM8 or GFP-MCM9 rescues and reduces the FANCD2 foci in both situations (10.2 ± 1.0 or 6.7 ± 0.5, respectively) (**Fig. 4C**). The quantification of FANCD2 foci does not differ between 8KO or 9KO cells in either NT or +MMC conditions consistent with MCM8/9 acting as a complex during fork stalling mediation. Interestingly, the monoubiquitination of FANCD2 does not appear to be impacted in 8KO or 9KO cell lines compared to WT, either in nontreated or MMC treated asynchronous or double thymidine block S-phase synchronized cells (**Supplementary Figure S6**). A higher molecular weight band consistent with ub-FANCD2 is visualized in all situations and conditions.

**Figure 4.**
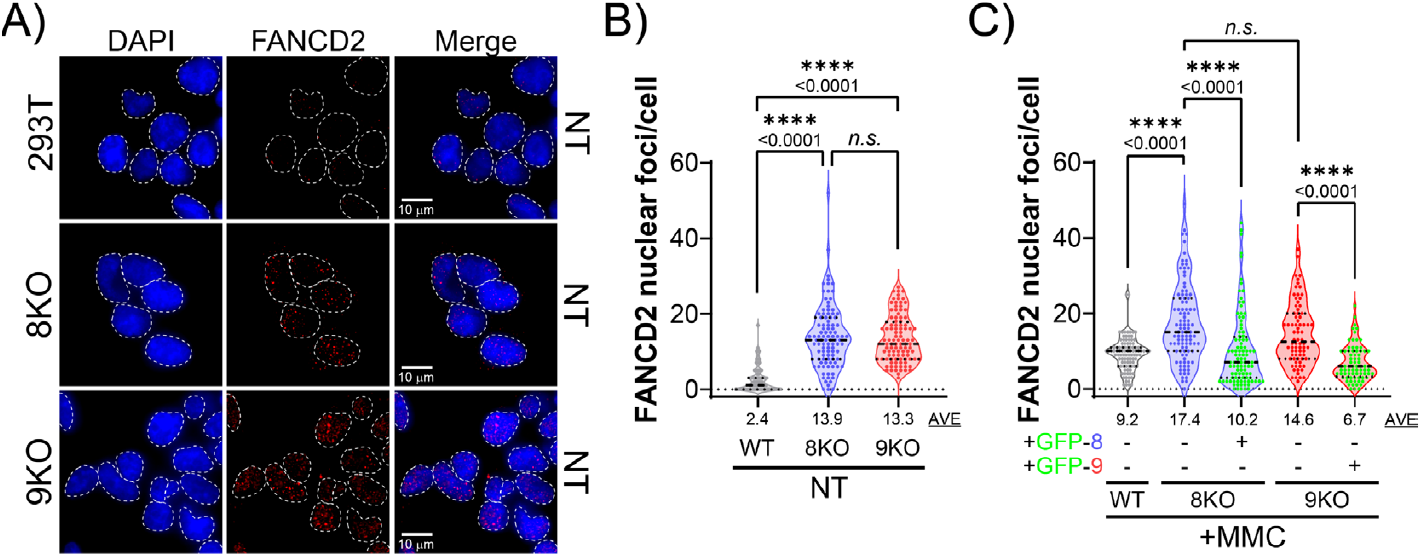
Knockout of MCM8 or MCM9 increases FANCD2 foci. A) Representative immunofluorescence images showing FANCD2 foci (red) in untreated WT, 8KO, or 9KO 293T cells. Cells are counterstained with DAPI (blue) to identify nuclei (white dashed areas). Quantification of FANCD2 foci per nucleus in B) untreated or C) MMC treated (3 μM for 6 h) WT, 8KO, or 9KO cells as well as 8KO and 9KO cells complemented with GFP-MCM8 or GFP-MCM9, respectively. Complementation rescues the elevated FANCD2 foci to basal levels. At least 100 nuclei were analyzed per condition and the averages for each experiment are indicated.

### 3.3 FANCD2 is required for the proper recruitment of MCM9

Next, we wanted to determine the dependency of FANCD2 in forming nuclear GFP-MCM9 foci with MMC damage. GFP-MCM9 was transfected into WT or FANCD2 knockout (FD2KO) 293T cells (**Supplementary Figure S7**) either nontreated or MMC treated, and the number of GFP-MCM9 nuclear foci were quantified per cell (**Fig. 5A-B**). In nontreated conditions, the number of nuclear GFP-MCM9 foci is low in WT cells (0.5 ± 0.1) and in FD2KO cells (1.8 ± 0.3) as expected from previous experiments. WT 293T cells treated with MMC show an increases number nuclear GFP-MCM9 foci as also seen in **Figure 1A**. However, FD2KO cells treated with MMC show a drastic reduction in GFP-MCM9 foci per cell nuclei (3.7 ± 0.4), consistent with MCM9 being dependent on FANCD2 for proper nuclear localization to affect a DNA damage response. Restoration of GFP-MCM9 foci occurred as expected upon the transfection of FD2-mCherry in both nontreated (0.5 ± 1) and MMC treated (13.8 ± 1.2) FD2KO cells (**Fig. 5C-D**). Colocalization of GFP-MCM9 and FD2-mCherry nuclear foci was also restored in FD2KO cells treated with MMC.

**Figure 5.**
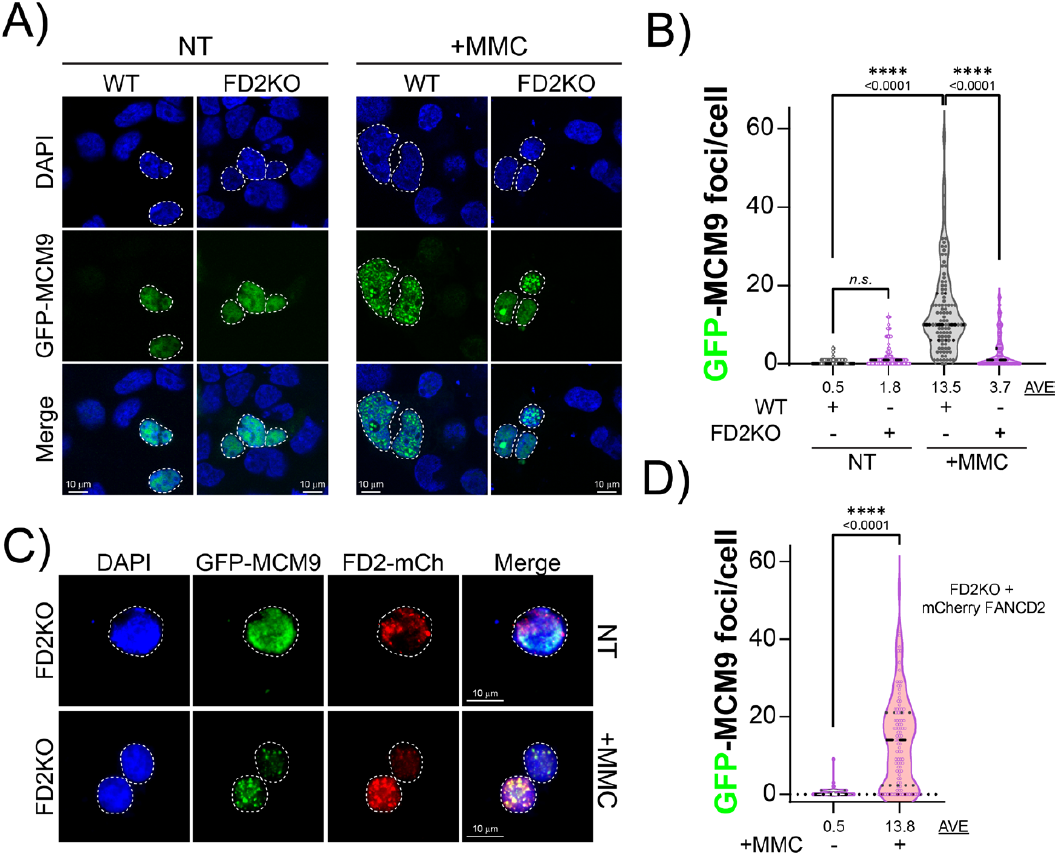
FANCD2 is required for the proper recruitment of MCM9. A) Representative immunofluorescence images showing GFP-MCM9 in untreated or MMC treated (3 µM for 6 h) WT or FD2KO cells. B) Quantification of GFP-MCM9 foci per nucleus in WT and FD2KO cells for untreated or MMC-treated conditions. C) Complementation of FD2KO cells with pRPmCherry-hFANCD2 followed by transfection with pEGFP-MCM9 restores GFP-MCM9 foci formation and colocalization with FD2-mCherry in MMC-treated cells. D) Quantification of MCM9 foci in FD2KO complemented cells ± MMC. At least 100 nuclei were analyzed per condition.

### 3.4 MCM9 is epistatic with FANCD2

FANCD2 and MCM9 are both involved replication stress response to protect and stabilize stalled forks and prevent DSBs. To test whether their roles are additive or synergistic when both FANCD2 or MCM9 are absent, we examined the frequency of DSBs measured by γH2A.X in FD2KO or 9KO cells with corresponding siRNA knockdown and treatment with MMC (**Fig. 6A**). In nontreated cells, the knockout of either FANCD2 or MCM9 resulted in a significant increase of γH2A.X foci (25 ± 2 vs. 25 ± 3) compared to the WT (17 ± 1) (**Fig. 6B-C**). The addition of either siMCM9 in FD2KO cells (26 ± 2) or siD2 in 9KO cells (28 ± 2) did not significantly change the amount of γH2A.X foci in the untreated conditions. When the corresponding cell lines were treated with MMC, the number of γH2A.X foci increased in all cell lines compared to the control siRNA (**Fig. 6D-E**). With MMC, knockout of FANCD2 or MCM9 increased γH2A.X foci (46 ± 3 vs. 32 ± 2) compared to WT cells (22 ± 2). However, the addition of either siMCM9 in FD2KO cells (43 ± 3) or siD2 in 9KO cells (33 ± 2) again did not significantly alter the amount of γH2A.X foci with MMC. Therefore, there is a codependence, such that removal of either FANCD2 or MCM9 results in increased DSBs but that removal of both does not have any additional effect, suggesting that MCM9 is epistatic with FANCD2, or within the same DNA damage response pathway.

**Figure 6.**
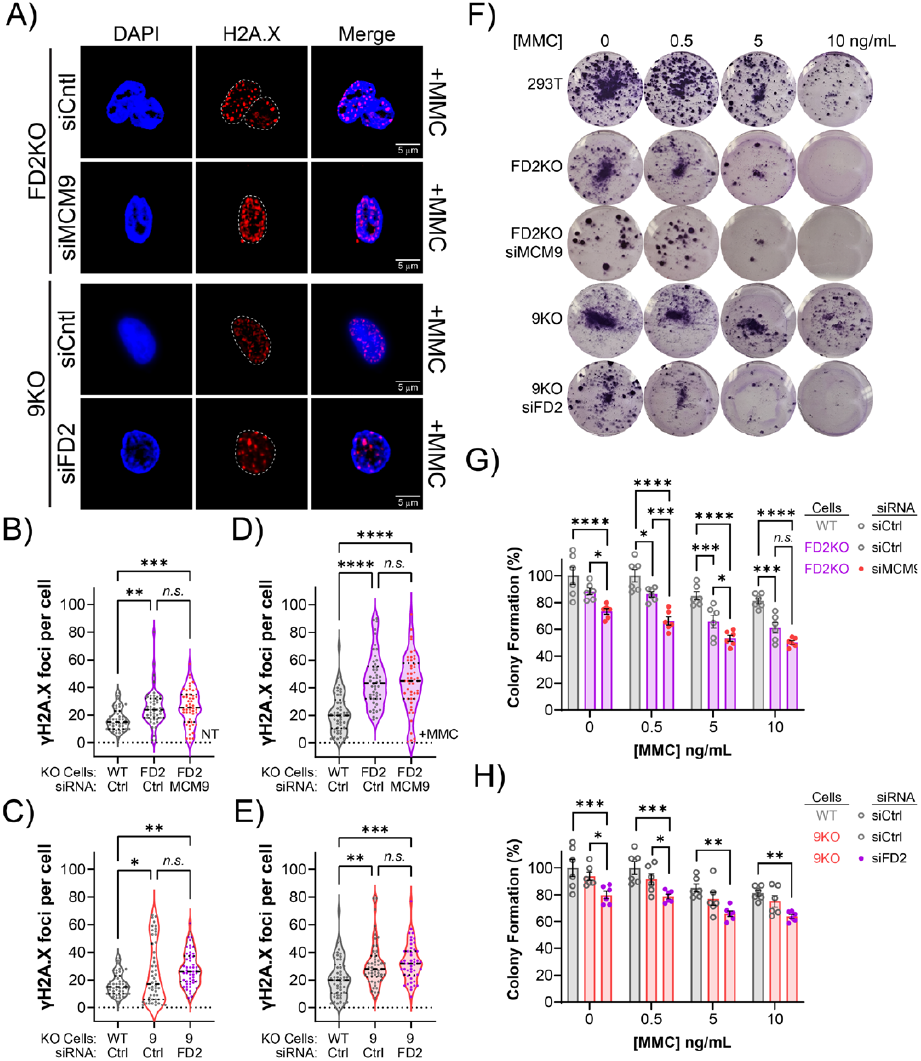
Knockout and co-depletion of FANCD2 and MCM9 does not increase DBS or reduce cell survival with MMC treatment. A) Representative immunofluorescence images showing γH2A.X foci (red) in 9KO or FD2KO cells with siCtrl or reciprocal siRNA knockdown treated with MMC (3 μM for 6 hours). Quantification of γH2A.X foci in B-C) nontreated or D-E) MMC treated conditions (n>50 cells). F) Representative crystal violet clonogenic survival assays images in 6-well plates seeded with WT, FD2KO, 9KO, FD2KO-siMCM9 co-depleted, or 9KO-siFD2 co-depleted cells treated with indicated concentrations of MMC for 24 hours. Colonies were stained after 12 days with crystal violet. Quantification of the colonies counted in G) FD2KO H) MCM9KO cells relative to the WT control.

To test whether the relationship between FANCD2 and MCM9 seen for DSBs also influences cell viability, we performed short-term cell survival assays using similar siRNA knockdowns in the reciprocal cell lines and treated them with increasing concentrations of MMC (**Supplementary Figure S8**). The relative cell viability was measured using CellTiter-Blue, which reports on the metabolic activity of cells compared to untreated controls. Both FD2KO and 9KO cell lines were moderately sensitive to increasing concentrations of MMC compared to WT cells; however, siFD2 or siMCM9 knockdown in the reciprocal knockout cells did not have any further decrease in viability. To be certain, we performed longer term colony formation assays that assessed cell proliferation over the same MMC concentrations (**Fig. 6F**). The siRNA depletion of MCM9 in FD2KO cells modestly decreased colony formation compared to FD2KO cells alone at 0.5 and 5 ng/mL concentrations but not at the higher 10 ng/mL concentration (**Fig. 6G**). Similarly, siFD2 in 9KO cells only showed decreased survival at 0.5 ng/mL but not at the higher concentrations of 5 or 10 ng/mL. Overall, the co-deletion/depletion of FANCD2 and MCM9 did not have any substantial additive or synergistic effect on cell survival when treated with MMC. Together, we conclude that MCM9 and FANCD2 act within the same replication stress induced DNA damage response pathway. MCM8/9 is epistatic with FANCD2 and functions downstream of FANCD2I to protected stalled forks from crosslink damage.

## 4. Discussion

DNA crosslinks produce toxic lesions that stall the replication fork and require repair before restarting replication. Crosslink repair is directed by the Fanconi anemia (FA) pathway which consists of at least 22 responding proteins. The key regulatory process for repair recognition involves the mUb of FANCD2 and FANCI by the FA core complex, which causes the complex to clamp around the duplex containing ICL at a fork junction (**Fig. 7**) [34-36]. Afterwards, ICL repair is proposed to take place through recruitment of the SLX4/XPF/ERCC1 incision complex to remove the crosslinked DNA [37, 38]. However, the steps between recognition and incision, translesion synthesis (TLS), and HR, are still ambiguous.

**Figure 7.**
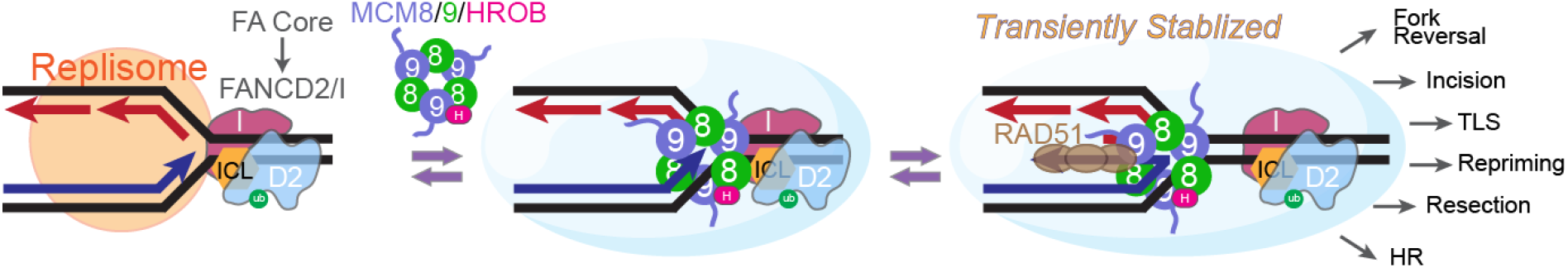
Model for FANCI/D2 recruitment of MCM8/9 at stalled forks. Upon sensing interstrand crosslink (ICL, *orange*) induced DNA damage, the Fanconi anemia (FA) core complex monoubiquinates (ub, *green*) the FANCD2-FANCI (D2I, *magenta & cyan*) complex. The ubD2-I complex binds to damaged DNA at the replication fork junction and recruits the MCM8/9/HROB (*green, blue & pink*) complex to the site. MCM8/9/HROB subsequently facilitates the destabilization of the leading strand to begin the reversed fork process by depositing RAD51 (*brown*). Our model suggests that FANCD2/I functions upstream of MCM8/9, acting prior to MCM8/9 assembly at damaged forks to act as an intermediary for downstream repair pathways.

The MCM8/9 complex also responds specifically to crosslink damage and acts to stabilize stalled forks from rampant fork reversal and nuclease degradation leading to increased DSBs, seemingly intersecting the FA pathway [13]. MCM8/9 acts upstream of RAD51 and BRCA1 (two other FA proteins) and also early in the crosslink repair process. Therefore, we sought to examine whether FANCD2/I and MCM8/9 overlap within the same crosslink repair pathway. To our surprise, we find that not only are FANCD2/I and MCM8/9 within the same pathway, but they also directly interact and respond equally to and colocalize completely with MMC damage. FANCD2/I is present directly upstream of MCM8/9 during crosslink damage recognition, where MCM9 is then epistatic with FANCD2. The only thing preventing us proposing that MCM8 and MCM9 are also Fanconi genes is that patients with those mutations do not have any history of anemia [23, 24] even though knockout *Mcm8*^*-/-*^ or *Mcm9*^*-/-*^ mice develop myeloid tumors that resemble myelodysplastic and anemia [39].

MCM8 and MCM9 both have core N-terminal DNA binding and AAA^+^ ATPase domains that form an alternating hexameric ring shaped helicase complex [14], but MCM9 also contains an extended and primarily unstructured C-terminus (CTE) that contains motifs required for formation of crosslink induced nuclear foci. Based on the co-IP mapping of the FANCD2 interaction with MCM8/9, the MCM9 CTE is unnecessary for maintaining this interaction. FANCD2 even interacts with an ATPase dead version of MCM8/9 or when we knock down the MCM8/9 activating protein, HROB, known to enhance unwinding. It is also interesting that knockdown of HROB prevents MCM9 nuclear foci formation with damage, suggesting that HROB serves as a loader, activator, or stabilizer for MCM8/9 at damage sites, consistent with other reports [19, 20].

However, the BRCv motif within the CTE of MCM9, is necessary for the stable localization of the MCM8/9 complex and to recruit RAD51 for complete fork stabilization when cells are treated with MMC [22]. Again, this BRCv motif is required for MMC induced MCM9 foci formation but not for the FANCD2 interaction. Furthermore, mutation of the BRCv motif does not affect FANCD2 foci formation with MMC, again suggesting a dependency for FANCD2 in localizing MCM8/9 and temporal organization. This is somewhat difficult to reconcile; however, it seems that although the core domain of MCM8/9 interacts with FANCD2 in all situations, stable loading of MCM8/9 leading to foci formation requires FANCD2, the active form of the MCM8/9 helicases (ATPase and HROB), and the ability to recruit RAD51 (BRCv motif) (**Fig. 7**).

Based on the clamp structure of FANCD2/I on DNA and its rapid translocation ability along the template to bounce off genomic roadblocks but stall at primer/template junctions [35], it is tempting to speculate that MCM8/9 is also located specifically at those sites. Stalled replication forks that encounter ICL damage will require FANCD2/I to identify and initiate a downstream repair process. Although it is thought that fork convergence will initiate ICL repair after removal of the CMG helicase [40, 41], it is also possible that single sided fork ICL encounters could initiate repair through fork reversal and stabilization, ICL incision, TLS, and restart through a variety of proposed mechanisms [42, 43]. FANCM plays an early role in recruiting the FA core complex through interactions with AND-1 at the front of the replisome [44]. It was shown previously, the restart from an ICL was dependent on the ATR kinase, DONSON, FANCD2, and FANCM [45]. However, ATR kinase activity is dependent on RPA association at stalled replication forks [46, 47], which requires more significant fork remodeling and RPA binding after the ICL encounter. Here, we propose that MCM8/9 is an intermediary in this ICL repair process, where it begins the fork remodeling process by interacting with ub-FANCD2/I to recruit and deposit RAD51 on single stranded regions of reversed forks created by its helicase activity. This would provide a temporal distinction for the ICL helicases where FANCM is important for mUb-FANCD2/I at the ICL to then recruit MCM8/9. It will be interesting to test the localization and codependency of FANCM and MCM8/9 in the future. MCM8/9 then initiates methodical fork remodeling by facilitating the deposition of RAD51 to stabilize stalled forks and activate ATR signaling. Afterwards, FANCD2/I and MCM8/9 will dissociate to allow for downstream effectors to carry out more fork reversal, resection, TLS, restart, or HR to finalize the repair process likely through different downstream pathways.

## CRediT authorship contribution statement

**Rashini Baragama Arachchi**: Conceptualization, Formal analysis, Investigation, Methodology, Validation Writing—original draft, Writing—review and editing; **Desmond Okafor**: Investigation and Formal analysis; **Andrew Snyder**: Investigation; **Michael Trakselis**: Conceptualization, Formal analysis, Visualization, Supervision, Funding acquisition, Writing—original draft, Writing— review and editing.

## Funding

Research reported in this publication was partially supported by the NIH (GM135791 and GM155805 to M.A.T.) and by Baylor University.

## Declaration of Competing Interest

The authors declare that they have no conflicts or competing interests with the contents of this article.

## Acknowledgements

We thank all members of the Trakselis laboratory for productive conversations and insights. We acknowledge the Baylor Molecular Bioscience Center (MBC) and the Center for Microscopy and Imaging (CMI) for providing instrumentation and resources aiding this project.

## Appendix A

Supplementary material

Supplementary data associated with this article including Table S1 and Figures S1-S8 can be found in the online version at ……

## Data Availability

The authors confirm that all other relevant data are present in the manuscript and its supplementary data. All biochemical data presented in this study including: value, gels, images, datasets as well as any strains or plasmids are available upon request to the corresponding author.

## Abbreviations

FA: Fanconi anemia
D2/I: FANCD2/FANCI
ICL: Interstrand crosslinks
HR: Homologous recombination
TLS: Translesion synthesis
MMC: Mitomycin C
HU: Hydroxyurea
siRNA: silencing RNA
KO: knockout
Co-IP: Co-immunoprecipitation
IP: Immunoprecipitation
BCA: Bicinchoninic acid assay
DMEM: Dulbecco’s Modified Eagle Medium
FBS: Fetal bovine serum; DNase I – Deoxyribonuclease

## Notes

### Competing Interest Statement

The authors have declared no competing interest.

