## Supplementary for "MCM8/9 and FANCD2 interact within a shared pathway in response to replication stress caused by DNA crosslinks"

**Supplementary Table S1: Oligonucleotides used in this study.**

**gRNA**

| **Name** | **Rest. Site** | **Sequence (5'-3')**[1] |
|---|---|---|
| gFD2-1 | *BsbI* | AACAGCCATGGATACACTTG |

**Primers for cloning**

| **Name** | **Rest. Site** | **Sequence (5'-3')** |
|---|---|---|
| MCM9F | *StuI* | AGGCCTAAAAAGAAGAGAAAGGTATAGATAACTGATCATAATCAGCCATACCACATTTG |
| 605R | *StuI* | CTATACCTTTCTCCTCCTTTTAGGCCTCTGCATTGAGGACTCCATGACTGACA |
| 648R | *StuI* | CTATACCTTTCTCTTCTTTTTAGGCCTCTCTTCACTCAAGAGGCTCTGCAG |
| 800R | *StuI* | CTATACCTTTCTCTTCTTTTTAGGCCTAAGCAACCCAATGTCCACTTTGCTCC |

**siRNA sequences**

| **Name** | **Company** | **Catalog#** |
|---|---|---|
| MCM9 | Sigma | SASI_Hs01_00041744, SASI_Hs01_00041746 |
| FANCD2 | Sigma | SASI_Hs01_00137854; SASI_Hs02_00307032, SASI_Hs01_00137857 |
| HROB | Sigma | SASI_Hs02_00356457; SASI_Hs01_00035908 |

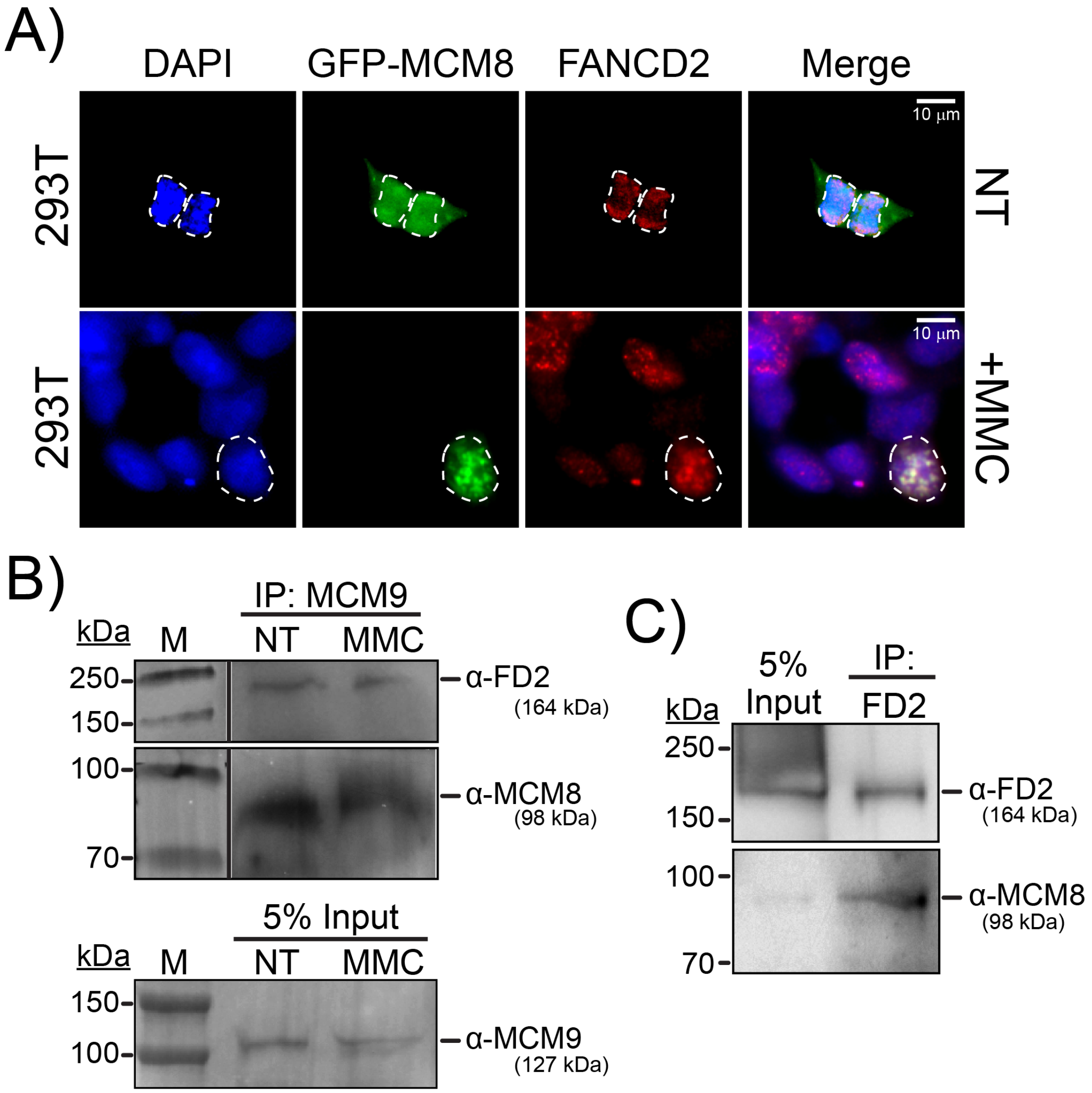


**Supplementary Figure S1. MCM8 and MCM9 interact with FANCD2.** A) Immunofluorescence of 293T cells transfected with pEGFP-MCM8 either nontreated (NT) or treated with 3 μM MMC (+MMC) for 6 hours. Endogenous FANCD2 (red) was stained with α-mouse FANCD2 antibody and GFP-transfected cell nuclei are outlined (white dashed circle). B) Endogenous Co-IP of MCM9 with FANCD2 and MCM8. Bottom panel is the 5% input of whole cell lysates probing for MCM9. C) Reverse Co-IP of FANCD2 probing for interaction with MCM8. Left lane is 5% input of whole cell lysate.

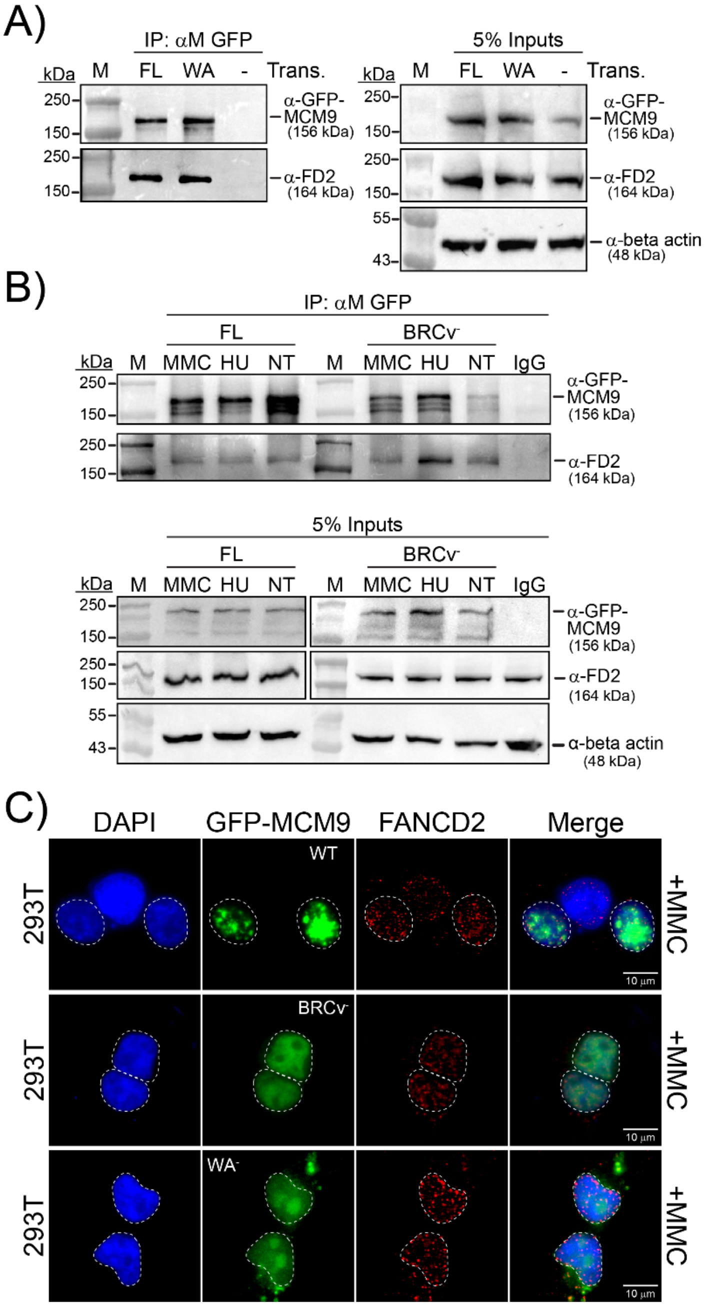


**Supplementary Figure S2: Mutation of either the Walker A or BRCv motifs do not disrupt the interaction with FANCD2.** A) Co-IP of GPF-MCM9 (FL or WA mutant) with FANCD2. 293T cells were transfected with pEGFP-MCM9 constructs or untransfected (- control) for 48 hours before treating with 0.5 μM MMC overnight. Whole cell lysates were IP'd with ChromoTek GFP-trap beads. Elutes were resolved by SDS-PAGE and immunoblotted for the indicated proteins. Left panel: immunoprecipitated proteins, right panel: 5% inputs. Molecular weight markers are indicated on the left and the position of constructs on the right with indicated theoretical molecular weights (kDa). B) FANCD2 interacts with BRCv mutant of MCM9. Cells were transfected with GFP-MCM9 (FL or BRCv mutant) and treated as in A) or with 2 mM HU overnight. Top panel: immunoprecipitated proteins, bottom panel: 5% inputs. C) WT 293T cells were transfected with pEGF-MCM9 (WT or BRCv or WA mutants) (green) and treated with 3 μM MMC (+MMC) for 6 hours. Endogenous FANCD2 (red) was stained with α-FANCD2 antibody.

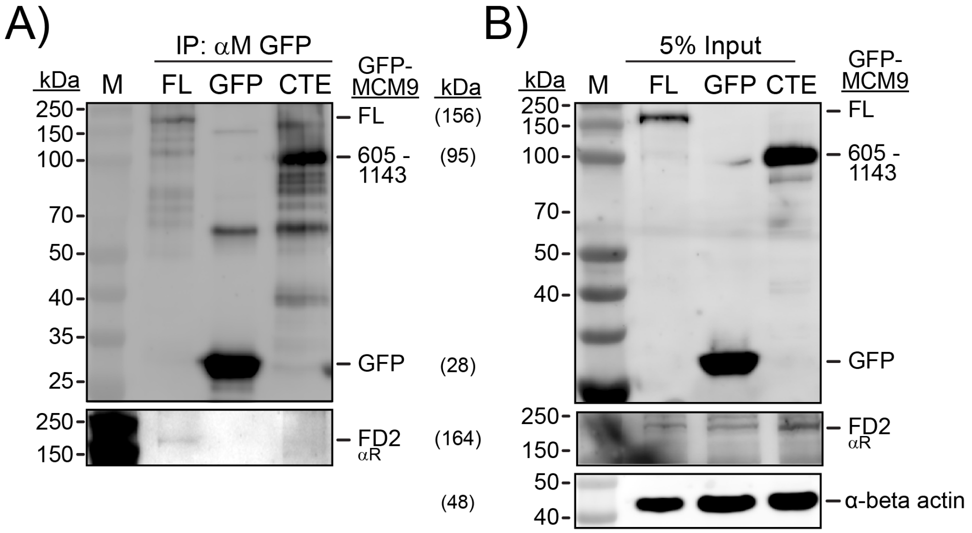


**Supplementary Figure S3: Control Co-IP with GFP or CTE of MCM9 do not interact with FANCD2.** A) 293T cells were transfected with pEGFP-MCM9$^{FL}$, pEGFP (empty), or pEGFP-MCM9$^{605\text{-}1143}$ for 48 hours. Whole cell lysates were IP'd with ChromoTek GFP-trap beads. Elutes were resolved by SDS-PAGE and immunoblotted for the indicated proteins. B) 5% Input blots of the protein lysates. Molecular weight markers are indicated on the left and the position of constructs on the right with theoretical molecular weights (kDa).

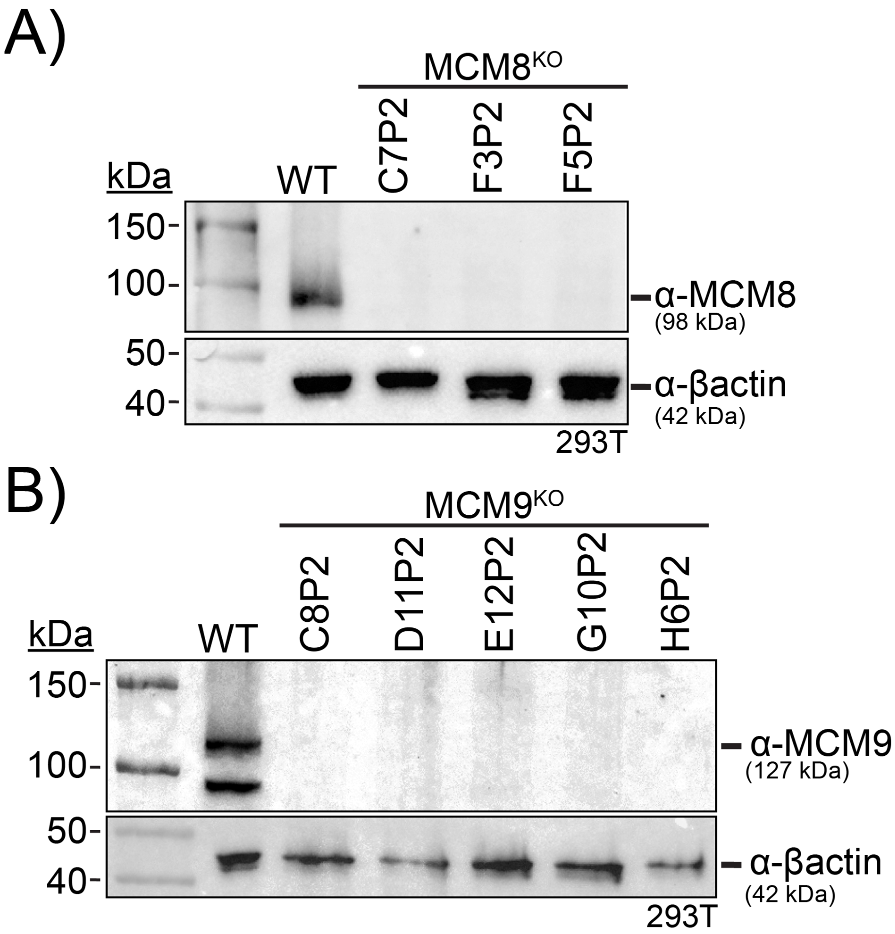


**Supporting Figure S4: Confirmation of knockout MCM8$^{KO}$ and MCM9$^{KO}$ 293T cell lines.** Whole cell lysates of selected clones were probed for the absence of A) MCM8 or B) MCM9. β-actin was used as a loading control in all lysates.

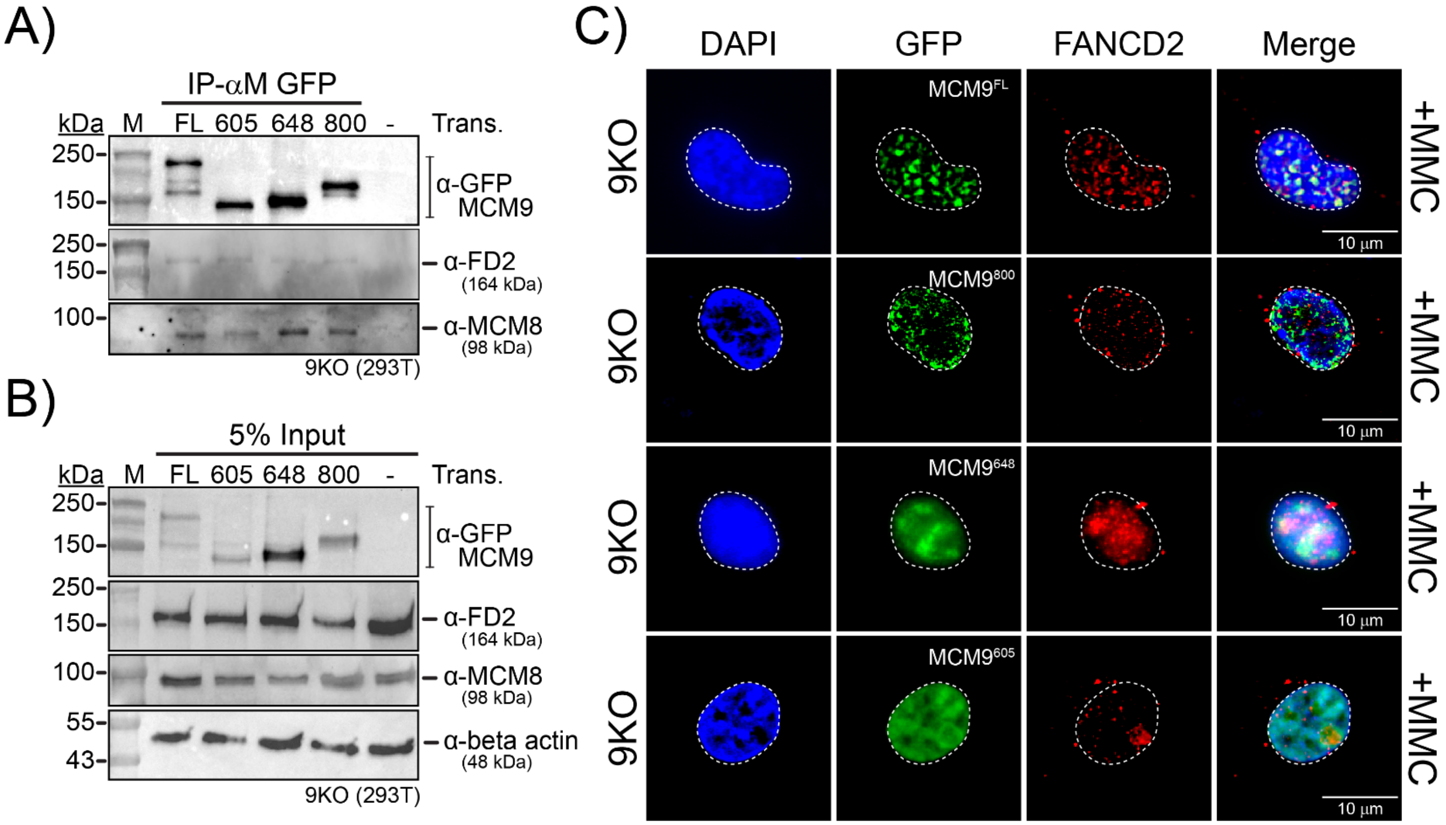


**Supporting Figure S5: Transfection of GFP-MCM9 in MCM9KO cells restores FANCD2 interaction and colocalization.** A) 9KO 293T cells (C8P2) were transfected with pEGFP-MCM9$^{FL}$, MCM9$^{605}$, MCM9$^{648}$, MCM$^{800}$ or not transfected (control) for 48 hours before treating with 0.5 μM MMC overnight. Whole cell lysates were immunoprecipitated with ChromoTek GFP-trap beads. Whole cell lysates were IP'd with ChromoTek GFP-trap beads. Elutes were resolved by SDS-PAGE and immunoblotted for the indicated proteins. B) 5% Input blots of the immunoprecipitated protein lysates. Molecular weight markers are indicated on the left and the position of constructs on the right with theoretical molecular weights (kDa). C) 9KO 293T cells were transfected with pEGF-MCM9 protein constructs (green) and treated with 3 μM MMC for 6 hours or left untreated. Endogenous FANCD2 (red) was stained with α-mouse FANCD2 antibody. MCM9$^{FL}$ and MCM$^{800}$ shows foci (green) and colocalize with FANCD2 (red) while MCM9$^{648}$ and MCM9$^{605}$ did not form MCM9 foci but left the FANCD2 foci formation unaffected.

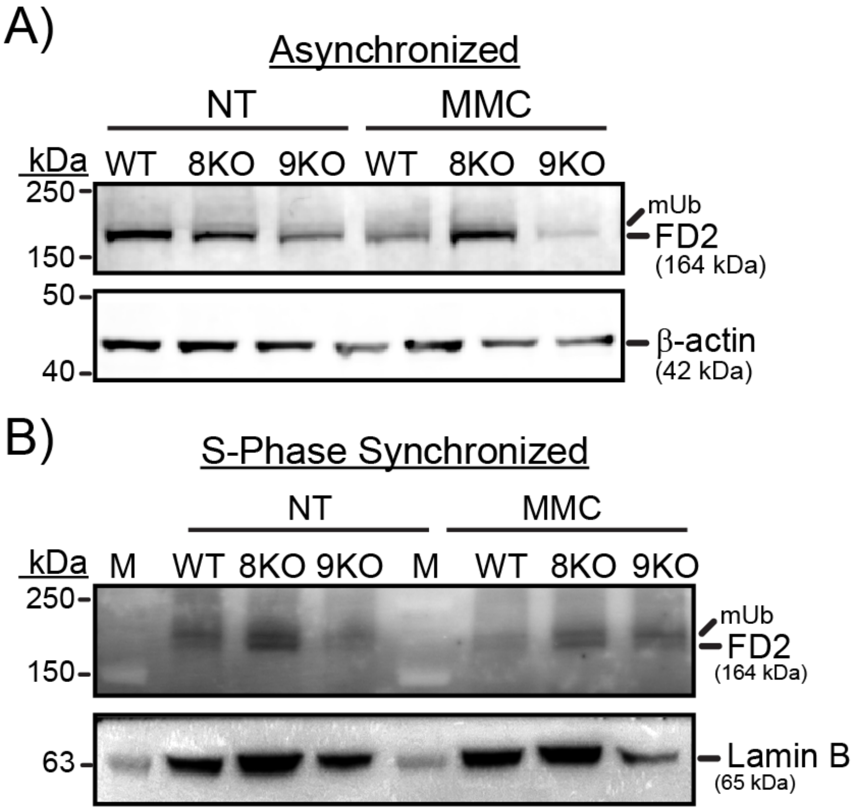


**Supporting Figure S6: Monoubiquitination (ub) of FANCD2 is unaffected in 8KO or 9KO cells treate with MMC.** A) WT, 8KO or 9KO (293T) cells were treated with 0.5 μM MMC overnight or left nontreated (NT). Whole cells lysates were resolved in 4-10% bis-tri gradient gels and probed with α-mouse FANCD2 or α-mouse beta actin (loading control) antibodies. B) Cells were synchronized to S-phase using a double thymidine block, treated with MMC or left nontreated (NT), and the whole cell lysates were resolved in 4-10% bis-tri gradient gels. α-mouse Lamin B was used as a loading control.

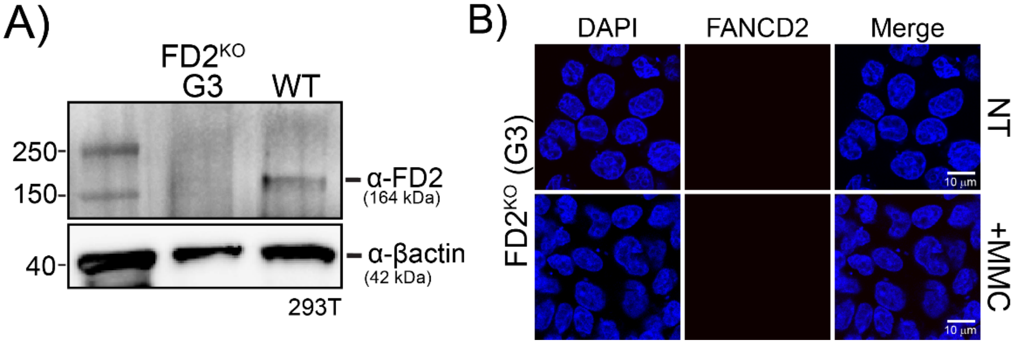


**Supporting Figure S7: Western blot and immunofluorescence validation of FANCD2$^{KO}$ cell strains.** A) Whole cell lysates of FD2KO (G3) and WT 293T cells were resolved in 10% SDS-gel and probed with α-mouse FANCD2 or α-mouse beta actin (loading control) antibodies. B) Representative immunofluorescence images showing the absence of signal for FANCD2 in MMC treated and nontreated (NT) FD2KO (G3) cells.

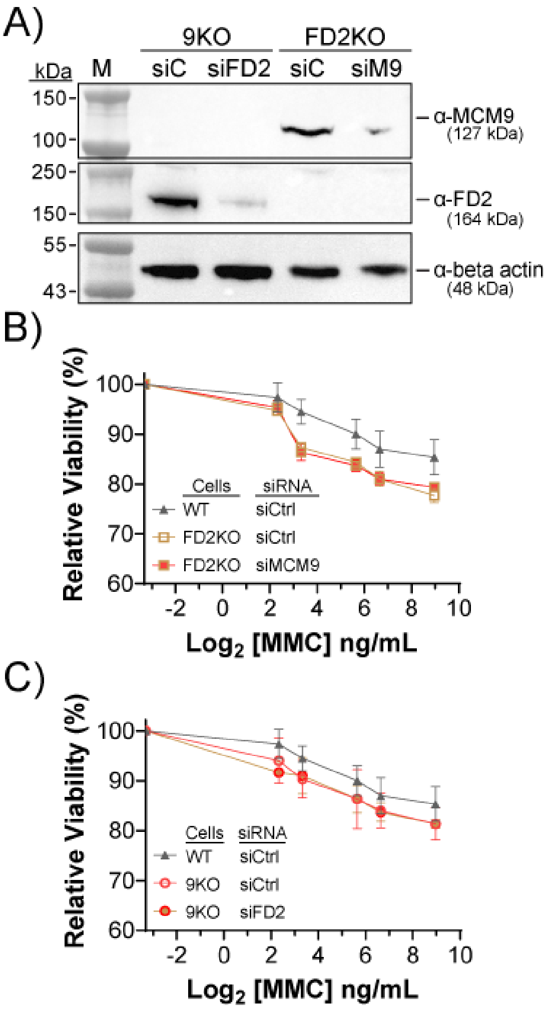


**Supporting Figure S8: Short term cell survival for codepletion and knockout of FANCD2 and MCM9 using CellTiter-Blue.** A) Western blot showing effective depletion of FANCD2 in $MCM9^{KO}$ cells and MCM9 depletion in $FANCD2^{KO}$ cells following siRNA transfection compared to control siRNA (siC) treated cells. Beta actin serves as the loading control. B) Relative cell viability of WT, FD2KO, and FD2KO + siMCM9 (co-depleted) following treatment with increasing concentrations of mitomycin C (MMC), as measured by CellTiter-Blue assay. C) Reciprocal experiment was performed in MCM9KO cells.
